# Molecular dissection of Class A PBP function uncovers novel features of the non-canonical *Clostridioides difficile* divisome complex

**DOI:** 10.1101/2025.05.29.656762

**Authors:** Gregory A. Harrison, Aimee Shen

**Affiliations:** Department of Molecular Biology and Microbiology, Tufts University School of Medicine, Boston, MA, USA

## Abstract

Cell division in bacteria is mediated by the “divisome,” a multiprotein complex that synthesizes the septal peptidoglycan needed to divide one cell into two. We recently showed that the major nosocomial pathogen *Clostridioides difficile* assembles a divisome that is fundamentally distinct from previously studied bacteria because it lacks functional orthologs of the septal peptidoglycan-synthesizing enzymes, FtsW and FtsI. While these enzymes were previously thought to mediate cell division in all walled bacteria, *C. difficile* instead uses the bifunctional Class A Penicillin Binding Protein PBP1 to mediate cell division. Here, we optimized a CRISPRi-based conditional expression system to define features within PBP1 that are critical for its essential functions. Our analyses identify a novel accessory domain that is required for PBP1 function and is conserved across Peptostreptococcaceae family PBP1 homologs. We further show that PBP1’s glycosyltransferase and transpeptidase activities are both strictly required for bacterial growth. While PBP1 glycosyltransferase activity is required for septum synthesis during cell division, PBP1’s transpeptidase activity is surprisingly dispensable for cell division, although TPase-deficient (PBP1^TPase*^) cells produce multiple aberrant septa. We demonstrate that the uncontrolled septum synthesis observed in PBP1^TPase*^ cells depends on the non-essential Class B PBP, PBP3, but PBP3’s catalytic activity is dispensable for this function. Since we also show that PBP3 is recruited to the divisome complex and forms a complex with PBP1, our analyses reveal a cryptic but important regulatory function for PBP3 in promoting *C. difficile* cell division.

**Author Summary:** Bacterial cell division is an ancient and essential process, but our molecular understanding of this process is primarily based on studies in a select few model systems. Recent work found that the major nosocomial pathogen *Clostridioides difficile* divides by a fundamentally distinct mechanism because homologs of the canonical septum synthesis complex used by virtually all walled bacteria are either missing from the *C. difficile* genome or have lost their function during vegetative division. This prior work revealed that septum synthesis in *C. difficile* is instead driven by the catalytic activity of the Class A Penicillin Binding Protein called PBP1, but the features of PBP1 that enable it to carry out this unusual function were unknown. In the current study, we perform detailed structure-function analyses to determine how *C. difficile* PBP1 has become specialized for this role. We find that PBP1 carries an unusual regulatory domain that is critical for its function. Our analyses also uncover an unexpected function for the non-essential enzyme PBP3 in cell division, identifying a new component of the *C. difficile* cell division complex. These analyses provide new insight into how bacteria can repurpose cell wall synthesis enzymes to fulfill essential functions in novel ways.

## Introduction

The broadly conserved process of bacterial cell division is mediated by a multi-protein complex called the “divisome.” The divisome is comprised of (i) a cytoskeletal scaffold based around the tubulin-like protein FtsZ that organizes the complex in a ring-like structure to mark the site of division and (ii) a transmembrane complex that includes the septal peptidoglycan synthases that drive cytokinesis (**Fig 1**). Decades of study in model systems have established that the septal peptidoglycan, which ultimately bisects the cell into two daughter cells, is generated by the transmembrane peptidoglycan synthase complex, FtsW-FtsI [1–3]. FtsW is a SEDS family glycosyltransferase that polymerizes the strands of peptidoglycan, and FtsI is a Class B Penicillin Binding Protein (bPBP) that crosslinks strands together through oligopeptide bridges. The activity and localization of the essential FtsW-FtsI complex is further controlled by the FtsQ-FtsL-FtsB transmembrane regulatory sub-complex. This sub-complex guides FtsW-FtsI to the FtsZ-ring at mid-cell and induces the septal PG synthase activity of FtsW-FtsI at the site of division [4–8]. The genes encoding these key divisome proteins are considered universally conserved across bacteria and most can be traced back to the last bacterial common ancestor billions of years ago [9].

**Figure 1.**
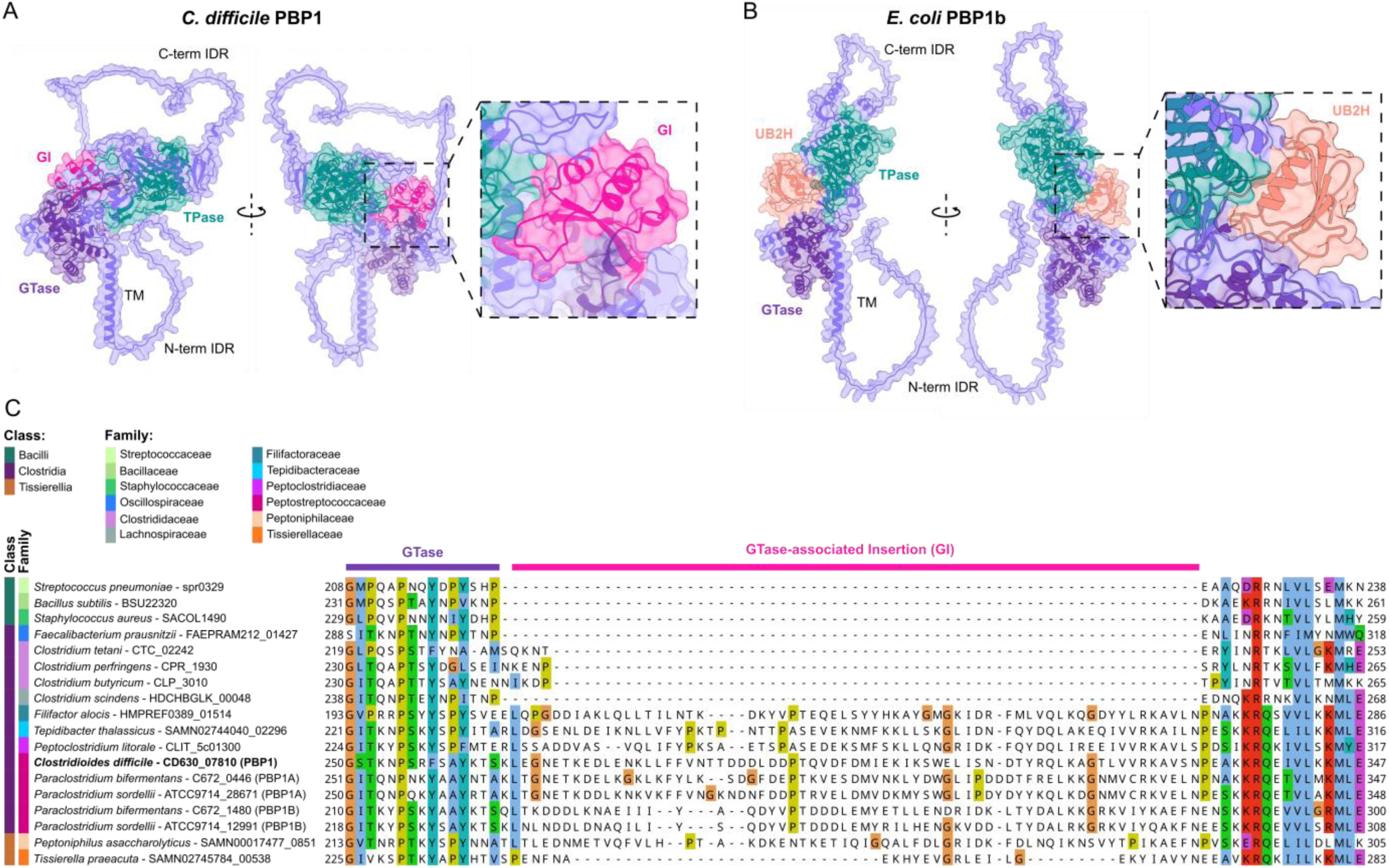
*C. difficile* PBP1 harbors an accessory domain conserved in evolutionarily related species as well as N- and C-terminal IDRs. (A-B) AlphaFold3 modeling of *C. difficile* PBP1 (A) and *E. coli* PBP1B (B) shows the C-terminal (C-term) and N-terminal (N-term) IDRs, as well as globular GTase-associated insertion (GI) and UB2H domains. TM indicates the transmembrane helix. (C) Protein sequence alignment of PBP1 homologs across Firmicutes species in the Bacilli, Clostridia, and Tissierellia classes was generated by Clustal Omega and visualized in Jalview. The class and family assigned to each species is based on current assignments in the NCBI taxonomy database [84].

While the FtsW-FtsI synthase complex was previously considered universally required for septal peptidoglycan synthesis in all walled bacteria [1,9–11], we recently discovered that *C. difficile* lacks homologs of FtsW and FtsI that function during vegetative cell division [12]. Furthermore, in contrast with previously studied bacteria [8,13–15], the *C. difficile* homologs of the FtsW-FtsI regulators FtsQ, FtsL, and FtsB are dispensable for vegetative cell division [12]. Therefore, *C. difficile* divides without the canonical septal PG synthase complex considered critical for division in virtually all other bacteria.

Since key canonical divisome proteins are either missing from the *C. difficile* genome (FtsW, FtsI) or are dispensable for division (FtsQ, FtsL, FtsB), *C. difficile* must use a unique suite of divisome proteins to carry out cell division [12]. Indeed, in lieu of the canonical FtsW-FtsI PG synthase complex, we recently showed that *C. difficile* uses the bifunctional Class A Penicillin-Binding Protein (**aPBP**) PBP1 to synthesize septal PG [12]. These analyses revealed a novel function for this class of enzymes in driving PG synthesis during cell division because aPBPs function primarily to reinforce the cell wall by contributing to bulk peptidoglycan biosynthesis in most well-studied bacteria [16,17]. In these systems, aPBPs play a supportive role to the SEDS-bPBP pairs that drive cell elongation (RodA-PBP2) and division (FtsW-FtsI), with aPBPs reinforcing the basement layers of cell wall generated by SEDS-bPBP pairs by filling gaps in the peptidoglycan and repairing damage [18–21]. Notably, in several Firmicutes species, this function is not essential for growth because aPBPs are completely dispensable for cell viability [22–24].

In contrast to this prevailing model of aPBP function, aPBPs play important roles in driving cell morphogenesis in certain polar-growing bacteria. For instance, Rhizobiales species such as *Agrobacterium tumefaciens* lack RodA-PBP2 altogether and instead rely on aPBP activity to drive cell elongation at cell poles [25,26]. Furthermore, in *Corynebacterium* and *Mycobacterium* species, RodA-PBP2 are dispensable for growth because aPBPs can mediate cell elongation in these organisms [27–30]. Thus, aPBP function is species-specific.

Our finding that *C. difficile* uses the aPBP PBP1 to drive cell division represents the first example, to our knowledge, of an aPBP driving cell division in the absence of the canonical FtsW-FtsI complex. Yet, how PBP1 has become specialized for this essential function is unclear. aPBPs in some species harbor accessory domains that contribute to their function at specific sites within the cell. For instance, in *E. coli*, the enzyme PBP1b harbors a UB2H domain, which is recognized by the outer membrane-bound lipoprotein LpoB through gaps in the peptidoglycan mesh [31–34]. LpoB activates PBP1b at these gaps, promoting PBP1b’s ability to fortify sites where the existing peptidoglycan layer is compromised [31,32,34]. While *E. coli* PBP1b is dispensable for growth and division in standard laboratory conditions, it localizes to the site of division and is thought to specifically reinforce the septal peptidoglycan that is generated by FtsW-FtsI [18,35–38]. Additionally, the *B. subtilis* aPBP PBP1 harbors an extracellular intrinsically disordered region (IDR) that allows it to sense gaps in the peptidoglycan and induce their repair [39], although *B. subtilis* PBP1 is dispensable for growth in standard laboratory conditions [22,40].

In this study, we investigate the function of specific PBP1 domains in *C. difficile* by developing a CRISPR-interference (CRISPRi)-compatible trans-complementation system to conditionally express mutant variants of *pbp1* in *C. difficile*. Using this system, we show that PBP1’s cytosolic and extracellular IDRs are dispensable for its function and identify a unique, novel regulatory domain of aPBPs that is critical for *C. difficile* PBP1 function and conserved in a subset of bacterial families. Our analyses of the role of PBP1’s catalytic activities reveal an unexpected role for the non-essential, monofunctional bPBP PBP3 in promoting septum synthesis. Thus, by using newly developed genetic tools in *C. difficile* to rapidly perform structure-function analysis of *C. difficile*’s unusual cell division machinery, we have defined key features of *C. difficile*’s essential Class A PBP and provided new insight into PBP function in bacteria.

## Results

### Unique structural features of *C. difficile* PBP1

Since *C. difficile* assembles a non-canonical divisome whose activity depends on PBP1, we sought to identify structural features that contribute to PBP1’s unique and essential divisome function. Alphafold3 [41] structural modeling predicts that PBP1 harbors two long intrinsically disordered regions (IDRs) at its cytosolic N-terminus and extracellular C-terminus (Fig 1A). The cytosolic N-terminal IDR is 46 amino acids long, and the extracellular C-terminal IDR is 108 amino acids long.

By comparing the structure of *C. difficile* PBP1 to well-studied PBP1 homologs in Firmicutes species such as *B. subtilis*, we identified a 66 amino acid domain insertion directly adjacent to *C. difficile* PBP1’s GTase domain that is missing from the PBP1 homologs in *B. subtilis* and other model Firmicutes species (**Fig. 1A**). This insertion domain, which we designated as the GTase-associated Insertion (GI) domain, is predicted to form a small globular fold near the linker region that connects the GTase and TPase domains (**Fig. 1A**). Intriguingly, the location of this domain is reminiscent of the UB2H domain of *E. coli* PBP1b (**Fig. 1B**). However, comparison of the GI domain of *C. difficile* PBP1 to the UB2H domain of *E. coli* PBP1b predicts that the GI domain is structurally distinct from the UB2H domain (**Fig 1A-B**).

By running FoldSeek [42] on the predicted structure of the GI domain, we found that this domain was only identified in aPBPs from a select few bacterial families, including the Peptostreptococcaceae, Peptoclostridiaceae, Tepidibacteraceae, Filifactoraceae, and Peptoniphilaceae. One eukaryotic protein was found to harbor a GI domain, an aPBP protein encoded in the genome of the protozoan parasite *Trichomonas vaginalis*. This protein is evolutionarily related to bacterial aPBPs and is encoded in a bacterial gene cassette recently acquired from a *Peptoniphilus* species through horizontal gene transfer [43]. Thus, aside from this instance of interkingdom horizonal gene transfer, the GI domain appears to be unique to bacterial aPBPs within a subset of bacterial families in the Firmicutes phylum.

Species within these five bacterial families primarily consist of Gram-positive (monoderm) anaerobic bacteria and occupy a range of anaerobic niches including the gut, oral cavity, vaginal tract, marine mud, and deep-sea hydrothermal vents [44–47]. Notably, species from these bacterial families were formerly classified under the Peptostreptococcaceae [46–48], suggesting that they may have shared features. The Peptostreptococcaceae, Peptoclostridiaceae, Tepidibacteraceae, and Filifactoraceae families are part of the class Clostridia and are primarily comprised of rod-shaped, Gram-positive bacterial species, many of which can form endospores [49]. In contrast, Peptoniphilaceae are non-spore-forming Gram-positive cocci of the class Tissierellia [50]. Aligning PBP1 homologs from representative Firmicutes species reveals that the GI domain is unique to this subset of bacterial families and not a general feature of aPBPs across the Firmicutes (**Fig. 1C**).

### *pbp1* conditional expression in *C. difficile* using CRISPRi-compatible complementation

We next sought to determine the functional significance of PBP1’s structural features, namely its GI domain and N- and C-terminal IDRs, in regulating PBP1 function in *C. difficile*. To rapidly probe the function of these domains in *C. difficile*, we developed a CRISPR-interference (CRISPRi) trans-complementation method to express conditional alleles of this essential gene. Specifically, we first knocked-down the expression of *pbp1* by integrating a xylose-inducible CRISPRi cassette [51] targeting *pbp1* into a neutral locus downstream of *pyrE* [52] (**Fig. 2A**, left). In the presence of xylose, this *pbp1*-knock down (KD) cassette represses the expression of the endogenous *pbp1* gene. We then complemented the *pbp1*-KD strain with a plasmid expressing CRISPRi-immune *pbp1* variants from an anhydrotetracycline (aTc)-inducible promoter [53] (**Fig. 2A**, right). The aTc-inducible *pbp1* constructs are immune to CRISPRi because they carry synonymous mutations in the sgRNA recognition sequence targeted by the CRISPRi cassette (*pbp1*_im_, **Fig. 2B**). Similar trans-complementation approaches have been used in *Borrelia burgdorferi* and *Mycobacterium tuberculosis* [54,55].

**Figure 2:**
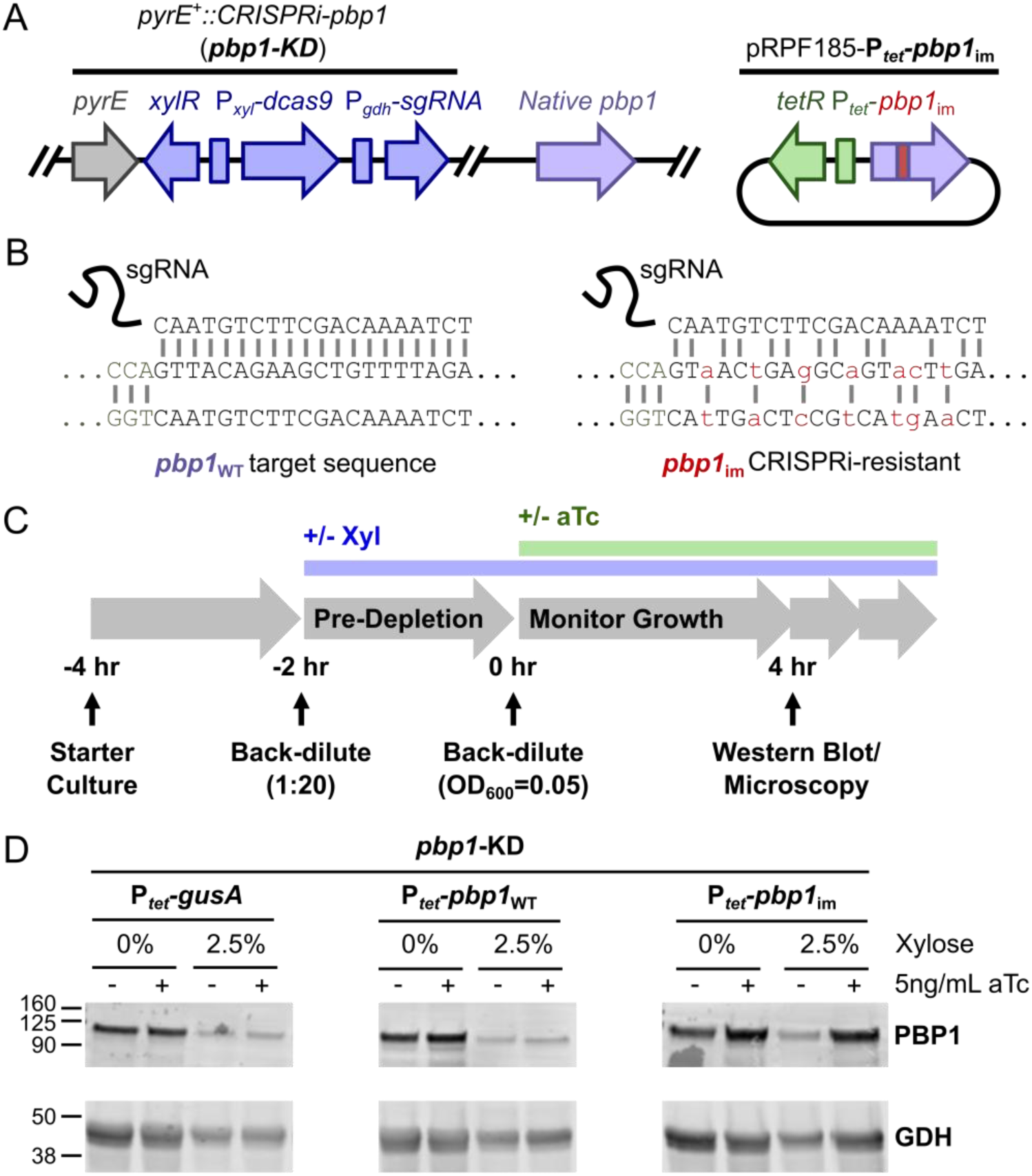
A CRISPRi trans-complementation system that enables the conditional expression of *pbp1* in *C. difficile*. (A) Schematic of the *pbp1*-KD strain. A xylose-inducible CRISPRi cassette targeting *pbp1* was inserted downstream of the *pyrE* locus in the *C. difficile* genome. aTc-inducible complementation constructs were supplied on the pRPF185 plasmid [53]. (B) To enable complementation of the CRISPRi-KD targeting *pbp1*, synonymous mutations were engineered into the sgRNA-targeting sequence in the plasmid to render *pbp1*_im_ resistant to CRISPRi targeting. (C) Schematic of the workflow for depleting PBP1 via the xylose-inducible CRISPRi-KD and rescuing PBP1 levels via the aTc-inducible expression of a CRISPR-immune *pbp1* variant carrying synonymous mutations (B). (D) Western blot analysis for PBP1 (96.5 kDa) and loading control GDH (46.0 kDa) using lysate from *pbp1*-KD *C. difficile* complemented with plasmids encoding *P_tet_*-*gusA*, -*pbp1*_WT_, or -*pbp1*_im_. PBP1 is depleted during growth in the presence of 2.5% xylose and restored only upon induction of *P_tet_*-*pbp1*_im_ with aTc. Blots are representative of three independent experiments, quantification of which can be found in Supplementary Fig. S1A.

To verify that our CRISPRi-compatible trans-complementation system complements *pbp1*-KD, we generated *pbp1*-KD *C. difficile* strains carrying plasmids expressing either (i) P*_tet_*-*gusA* (negative control), (ii) P*_tet_*-*pbp1*_WT_, which should remain susceptible to CRISPRi targeting, or (iii) P*_tet_*-*pbp1*_im_, which should be immune to CRISPRi targeting and rescue *pbp1* expression. We pre-depleted PBP1 by culturing each of these strains in the presence of 2.5% xylose for 2 hours to allow the endogenous levels of PBP1 to be diluted through ∼2-3 successive division cycles. We then exposed the bacteria to 0 or 5 ng/mL aTc for 4 hours to induce the expression of the complementation construct (**Fig. 2C**). With this experimental set-up, the bacteria experienced *pbp1-*KD for 6 hours (2 hours pre-depletion followed by 4 additional hours of treatment) and induction of the immune construct for 4 hours, all while maintaining the cultures in logarithmic growth (**Fig. 2C**). Exposure to xylose significantly decreased PBP1 levels, while aTc addition increased PBP1 levels in the P*_tet_*-*pbp1*_im_ strain but not the negative control strains (**Fig. 2D**)(Supplementary Fig. S1A). Notably, while the *pbp1-*KD did not fully deplete PBP1 (**Fig. 2D**)(Supplementary Fig. S1A), its levels were sufficiently reduced such that the growth inhibition and filamentation phenotypes associated with PBP1 depletion were observed (**Fig. 3**)[12,51]. Importantly, induction of the CRISPRi-immune *pbp1*_im_ using 5 ng/mL aTc fully complemented the growth inhibition caused by *pbp1*-KD, in contrast with the negative controls (*gusA* or *pbp1*_WT_, **Fig. 3A**, filled green diamonds). It should be noted that expression from the aTc-inducible promoter is leaky, such that even in the absence of aTc addition, we observed a modest restoration of growth in cells harboring P*_tet_-pbp1*_im_ (**Fig. 3A**; filled blue squares). Taken together, these analyses demonstrate that a CRISPRi trans-complementation system can be used to effectively knock-down native *pbp1* expression and heterologously express an ectopic, CRISPRi-resistant variant of *pbp1*.

**Figure 3:**
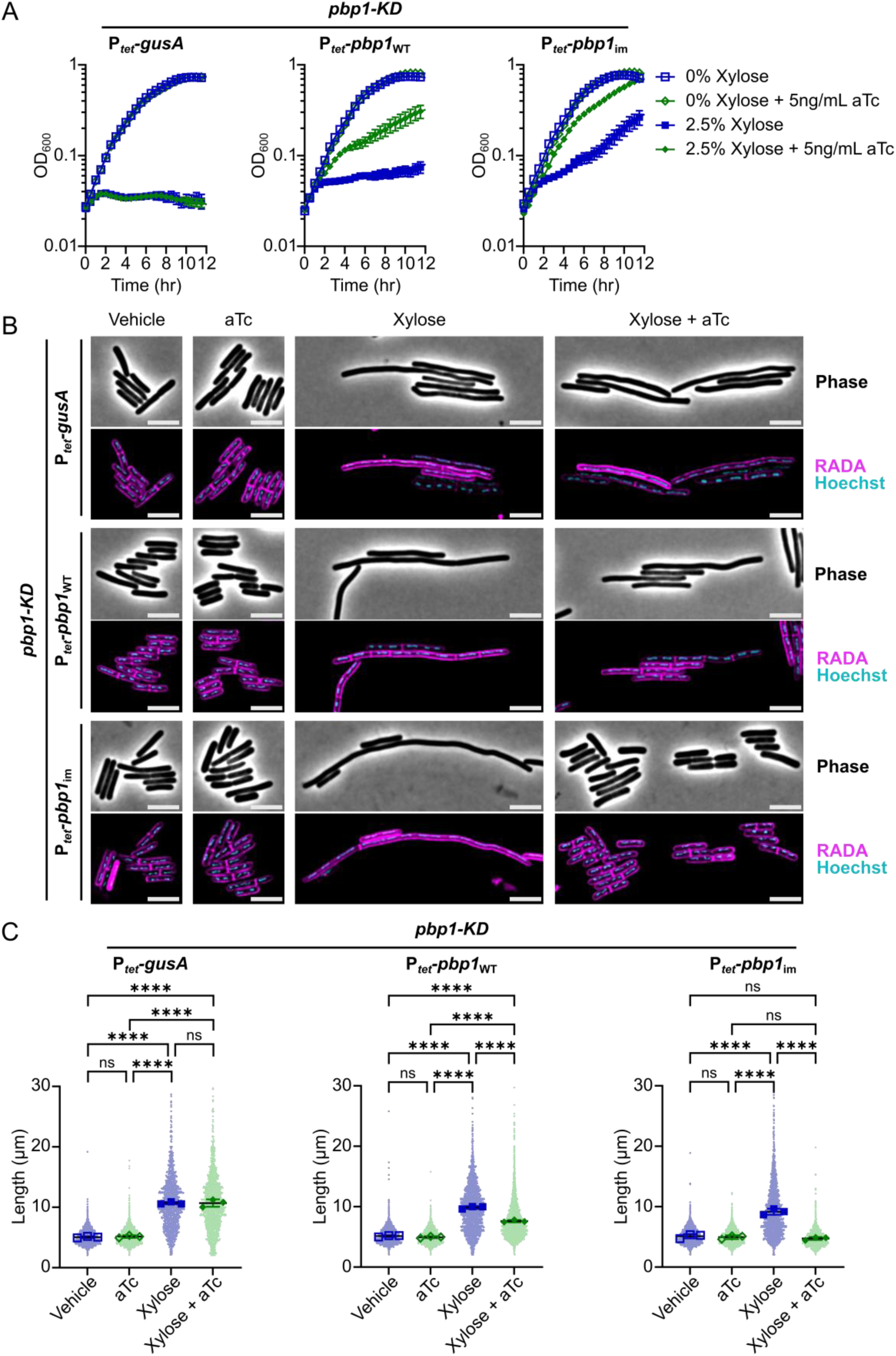
Rescue of *pbp1*-KD phenotypes using CRISPRi trans-complementation. (A) Growth of *C. difficile* strains containing the xylose-inducible *pbp1*-KD cassette and aTc-inducible *gusA*, *pbp1*_WT_, or *pbp1*_im_ complementation constructs was monitored by OD_600_. Mean and standard error were calculated across nine biological replicates. (B) Fluorescence microscopy analyses of cells exposed to vehicle or 2.5% xylose and/or 5 ng/mL aTc per the scheme in Fig. 2C. RADA labeling was used to visualize *de novo* peptidoglycan synthesis and Hoechst staining to visualize DNA. Images shown are representative of three independent experiments. Scale bars represent 5 μm. (C) Quantification of the length of >1200 cells across three independent experiments. Dots indicate individual cells, and the larger, outlined symbols represent the mean cell length from each replicate. The mean and standard deviation were calculated across replicates; statistical significance was determined by a one-way ANOVA with Tukey’s multiple comparisons test. ns, not significant; **** p<0.0001.

Since *pbp1*-KD also causes cell filamentation [12,51] due to defects in cell division (**Fig. 3B**), we examined the CRISPRi trans-complementation strains using microscopy. Specifically, we used the fluorescent D-amino acid RADA to visualize sites of active peptidoglycan synthesis and Hoechst to stain the bacterial nucleoid. As expected, *pbp1*-KD caused cell filamentation, although occasional septa could be detected within filaments, similar to previously published results [12,51]. Conditional expression of *pbp1*_im_ restored normal cell morphology and cell length (**Fig. 3B-C**). Simply over-expressing *pbp1* caused no change in cell length (**Fig. 3C**), although a subtle but significant increase in cell width from ∼0.8 μm to ∼0.9 μm was observed (Supplementary Fig. S1B). Since increased bacterial cell width has also been observed in *E. coli* or *B. subtilis* over-expressing aPBP enzymes that contribute to sidewall synthesis [19,56], these data support the conclusion that PBP1 is involved in sidewall synthesis as well as septal PG synthesis, consistent with its localization to both the sidewall and septum [12].

### PBP1’s IDRs are not strictly required for growth or division in *C. difficile*

Having developed a CRISPRi trans-complementation system to express mutant variants of *pbp1*_im_, we next tested the function of mutant *pbp1*_im_ alleles. Notably, our system enables *pbp1* mutant alleles to be rapidly tested within 2-3 days compared to the multiple weeks required to assess the function of chromosomally-encoded mutant alleles. We first examined the functional requirements for the N-terminal cytosolic IDR and C-terminal extracellular IDR by introducing plasmids encoding PBP1 variants lacking the 46-amino acid N-terminal IDR (P*_tet_*-*pbp1*_im_^ΔNT^), the 108-amino acid C-terminal IDR (P*_tet_*-*pbp1*_im_^ΔCT^), or both IDRs (P*_tet_*-*pbp1*_im_^ΔNTΔCT^) into the *pbp1*-KD strain. Following xylose-inducible PBP1 pre-depletion and aTc-inducible trans-complementation (**Fig 2C**), we found that conditional expression of *pbp1*_im_^ΔNT^, *pbp1*_im_^ΔCT^, or *pbp1*_im_^ΔNTΔCT^ rescued the growth inhibition caused by *pbp1*-KD as measured by OD_600_ over time (**Fig. 4A**). Thus, neither the N-terminal nor the C-terminal IDR are required for PBP1 to support the growth of *C. difficile* in standard laboratory conditions. Importantly, all three of these truncated protein variants were stably produced during knock-down of the native *pbp1* (**Fig. 4B**)(Supplementary Fig. S2A). Conditional expression of *pbp1* ^ΔNT^, *pbp1* ^ΔCT^, or *pbp1* ^ΔNTΔCT^ all restored WT cell length (**Fig. 4C**)(Supplementary Fig. S2B), indicating that the N- and C-terminal IDRs are dispensable for PBP1 function during *C. difficile* cell division. While conditional expression of both *pbp1* ^ΔCT^ and *pbp1* ^ΔNTΔCT^ appeared to increase cell curvature, with comma-shaped cells frequently observed in these two mutants (**Fig. 4C**), this apparent increase in cell curvature was not statistically significant (Supplementary Fig. S3). Thus, our data show that PBP1’s IDRs are not strictly required for growth or division in standard laboratory conditions, in contrast with *B. subtilis* PBP1, whose IDR is required for normal growth and cell morphology [39].

**Figure 4:**
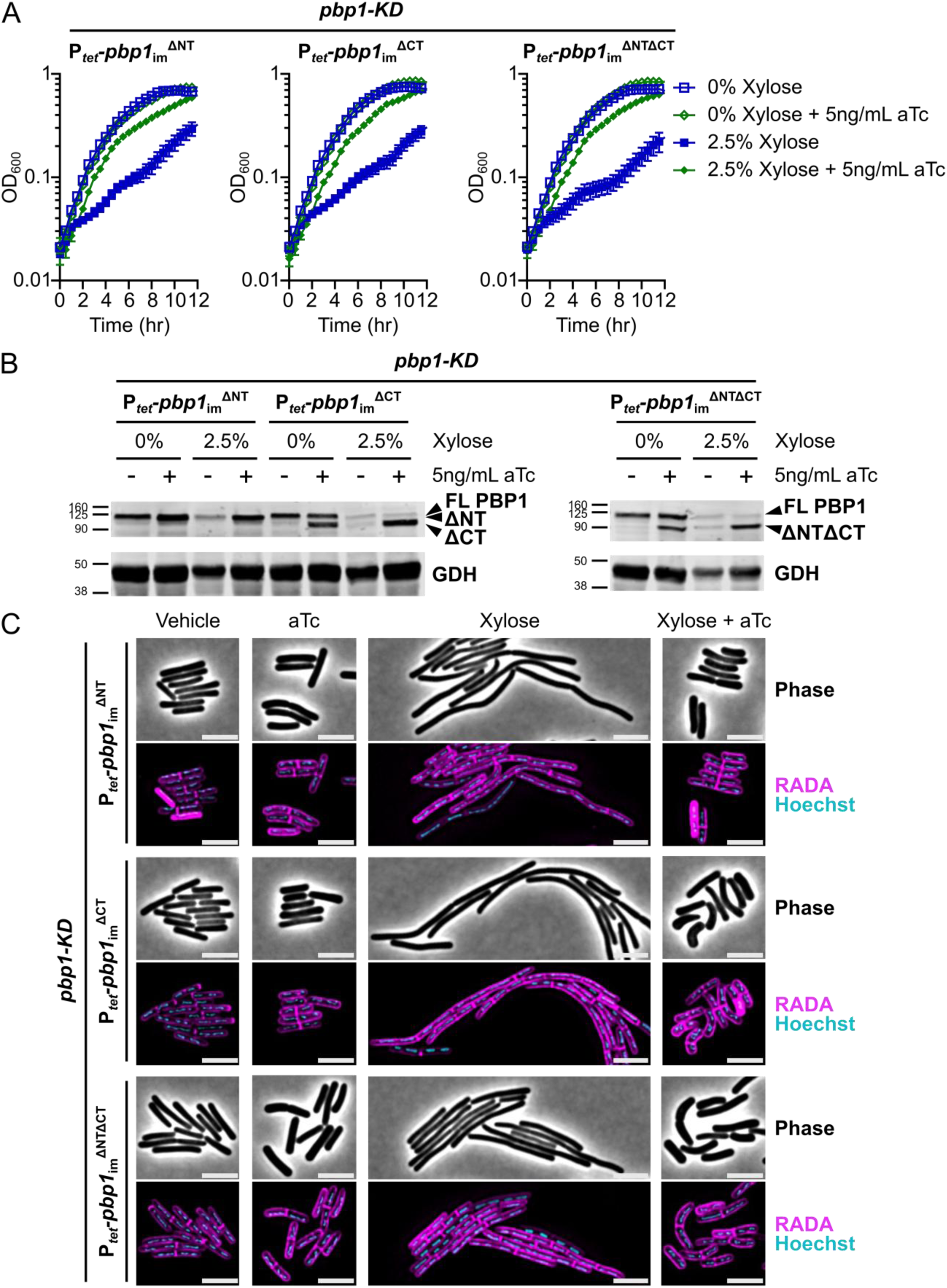
PBP1 N-terminal and C-terminal IDRs are not strictly required for growth or division. (A) Growth of *C. difficile* strains containing the xylose-inducible *pbp1*-KD cassette and aTc-inducible *pbp1*_im_^ΔNT^, *pbp1*_im_^ΔCT^, or *pbp1*_im_^ΔNTΔCT^ complementation constructs was monitored by OD_600_. Mean and standard error were calculated across nine biological replicates. (B) Western blot analyses of PBP1 in cells exposed to vehicle or 2.5% xylose and/or 5 ng/mL aTc per the scheme in Fig. 2C. GDH (46.0 kDa) was used as a load control. FL indicates Full-Length PBP1, and truncated protein variants are indicated. Blots are representative of three independent experiments, and their quantification is presented in Supplementary Fig. S2A. (C) Fluorescence microscopy analyses of cells exposed to vehicle or 2.5% xylose and/or 5 ng/mL aTc per the scheme in Fig. 2C. RADA labeling was used to visualize *de novo* peptidoglycan synthesis and Hoechst staining to visualize DNA. Dots indicate individual cells. Images shown are representative of three independent experiments. Scale bars represent 5 μm. Cell length quantification across replicates is presented in Supplementary Fig. S2B.

### The GI domain in PBP1 is critical for its function in *C. difficile*

We next assessed the functional significance of the other major structural feature in *C. difficile* PBP1 predicted by structural modeling, the GI domain, by introducing a plasmid encoding a GI domain mutant (PBP1^ΔGI^) into the *pbp1*-KD strain. The deletion construct lacks residues 267-328 and was designed based on AlphaFold3 modeling, which predicts that the functional domains of PBP1^ΔGI^ adopt a similar structure as in PBP1^WT^ (Supplementary Fig. S4A-B). Trans-complementation of the *pbp1*-KD with the aTc-inducible *pbp1* ^ΔGI^ construct revealed that loss of the GI domain prevented PBP1 function, as measured by bacterial growth (**Fig. 5A**). Importantly, PBP1^ΔGI^ was stably produced during our induction conditions based on western blot analyses (**Fig. 5B**)(Supplementary Fig. S5), indicating that the GI domain is critical for the proper functioning of PBP1.

**Figure 5:**
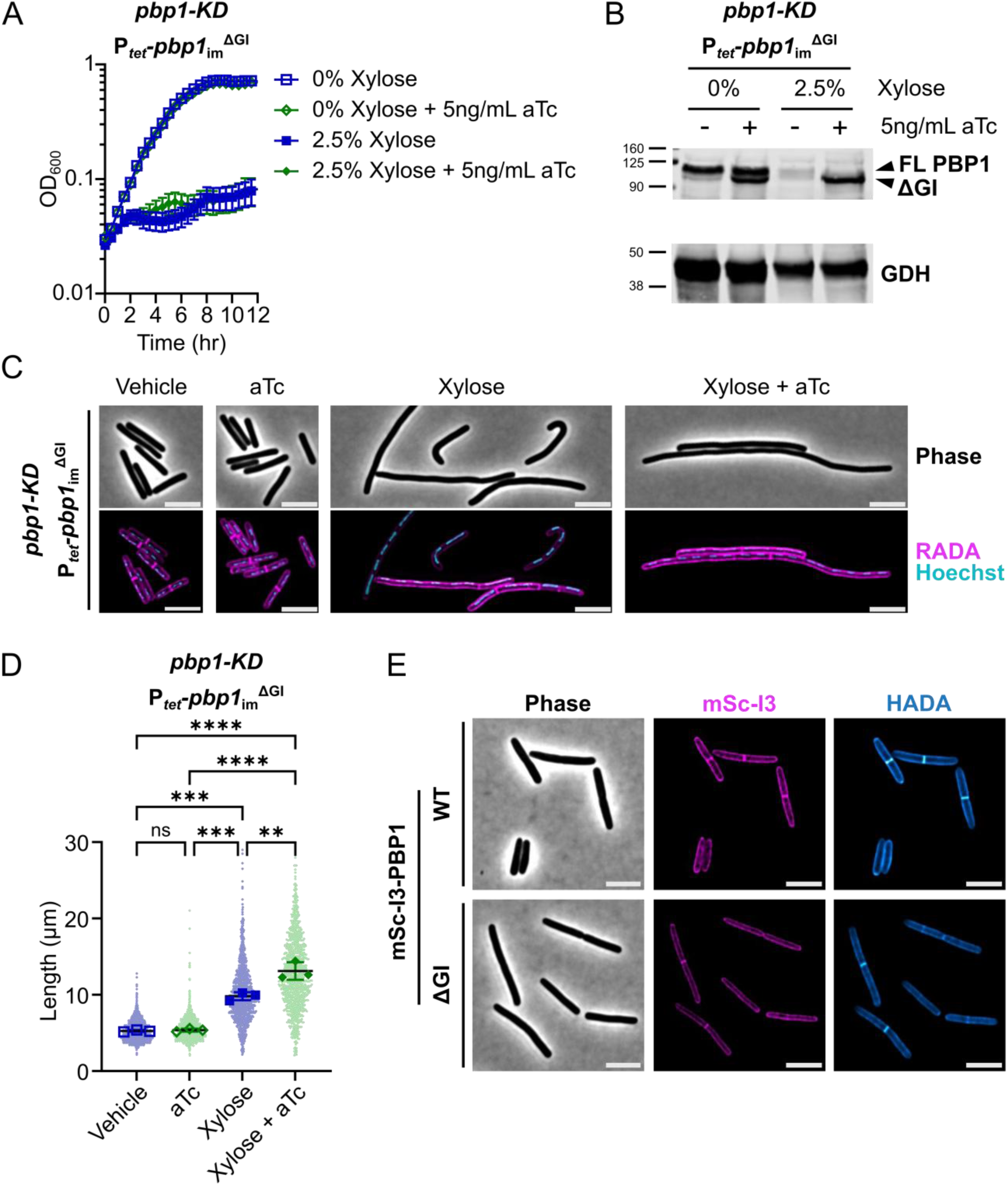
The GI domain is critical for PBP1 function in *C. difficile* growth and division. (A) Growth of the *C. difficile* strain containing the xylose-inducible *pbp1*-KD cassette and an aTc-inducible *pbp1*_im_^ΔGI^ complementation construct was monitored by OD_600_. Mean and standard error were calculated across nine biological replicates. (B) Western blot analyses of PBP1 in cells exposed to vehicle or 2.5% xylose and/or 5 ng/mL aTc per the scheme in Fig. 2C. GDH (46.0 kDa) was used as a load control. FL indicates full-length PBP1, and the band representing the ΔGI variant is indicated. Blots are representative of three independent experiments, and quantification across experiments is presented in Supplementary Fig. S5A. (C) Fluorescence microscopy of cells exposed to 2.5% xylose and/or 5 ng/mL aTc per the scheme in Fig. 2C. RADA labeling was used to visualize *de novo* peptidoglycan synthesis and Hoechst staining to visualize DNA. Images are representative of three independent experiments. (D) Quantification of cell length of >900 cells across three independent experiments. Dots indicate individual cells. The larger, outlined symbols represent the mean cell length from each replicate. The mean and standard deviation were calculated across replicates; statistical significance was determined between replicates by a one-way ANOVA with Tukey’s multiple comparisons test. ns, not significant; ** p<0.01, *** p<0.001, **** p<0.0001. (E) Localization of mScI3-PBP1 produced under the control of an aTc-inducible promoter using fluorescence microscopy. Cells were cultured in the presence of 5 ng/mL aTc for 1 hr, exposed to HADA for 10 min to label *de novo* peptidoglycan synthesis, and then fixed for microscopy. Scale bars represent 5 μm.

Consistent with the observed growth defects, *pbp1* ^ΔGI^ expression failed to reverse the filamentation phenotype caused by *pbp1*-KD, and even exacerbated its filamentation phenotype (**Fig. 5C,D**). Thus, the GI domain is required for PBP1 to synthesize septal peptidoglycan. To determine if the GI domain impacts the localization of PBP1 to sites of septal peptidoglycan synthesis, we analyzed the localization of PBP1 and PBP1^ΔGI^ variants with N-terminal fusions to mScarlet-I3 (mSc-I3)[57] in a WT background. The fusion protein constructs were engineered into an ectopic site in the chromosome, generating merodiploid strains that also express the native copy of *pbp1*. mSc-I3-PBP1^ΔGI^ localized to both the sidewall and at septa, similar to the mSc-I3-PBP1^WT^ [12](**Fig. 5E**). Thus, the GI domain is not required for proper protein localization and is instead likely required for the activity of PBP1 at the sites of peptidoglycan synthesis, either through direct allosteric activation of the GTase and/or TPase domains or by serving as an interaction site for unknown PBP1-regulatory proteins.

### PBP1 GTase and TPase activities are essential for growth

While these data reveal that the GI domain promotes PBP1 function, it is unclear whether PBP1’s distinct catalytic activities are specifically required during *C. difficile* growth and/or division. Given that inhibition of aPBP GTase activity with Moenomycin A (MoeA) recapitulates the effects of *pbp1*-KD [12], PBP1’s GTase activity is likely essential for its function in *C. difficile*. However, whether PBP1’s TPase activity is essential for its function remains unclear. Prior studies have proposed that PBP1’s TPase activity is dispensable for its function because β-lactam antibiotics that potently inhibit PBP1 TPase activity *in vitro* do not necessarily inhibit *C. difficile* growth [58]. Furthermore, the TPase activity of the essential division-specific bPBP FtsI is dispensable in both *B. subtilis* and *S. aureus* because other bPBPs can compensate for the loss of this activity [3,59]. In these species, the essential function of FtsI is to allosterically activate FtsW, while FtsI’s TPase activity can be supplied by functionally redundant TPases. We therefore tested whether *C. difficile* PBP1’s GTase or TPase catalytic activities are essential for its function using the CRISPRi conditional expression system.

To abrogate PBP1’s GTase activity, we generated a *pbp1*_im_ construct encoding two amino acid substitutions in the active site of the GTase domain within the conserved EDxxFxxHxG motif, E137Q/D138N (*pbp1* ^GTase*^). This mutation includes the catalytic glutamate residue and the adjacent aspartate, which is thought to play a key role in the catalytic mechanism [60]. We separately inactivated PBP1 TPase activity by generating a *pbp1*_im_ construct encoding a single amino acid change in the nucleophilic serine residue in the conserved SxxK motif within the TPase active site, S487A (*pbp1* ^TPase*^).

Conditional expression of *pbp1* ^GTase*^ and *pbp1* ^TPase*^ failed to complement the growth defect caused by *pbp1*-KD (**Fig. 6A-B**), indicating that the GTase and TPase activity of PBP1 are essential for *C. difficile* growth. These data further indicate that other TPases encoded by *C. difficile*, namely the two bPBPs and five L, D-TPases (LDTs) [12,61,62], cannot support the growth of *C. difficile* in the absence PBP1 TPase activity. Notably, both PBP1^GTase*^ and PBP1^TPase*^ were stably expressed in *C. difficile* as measured by western blot (Supplementary Fig. S6).

**Figure 6:**
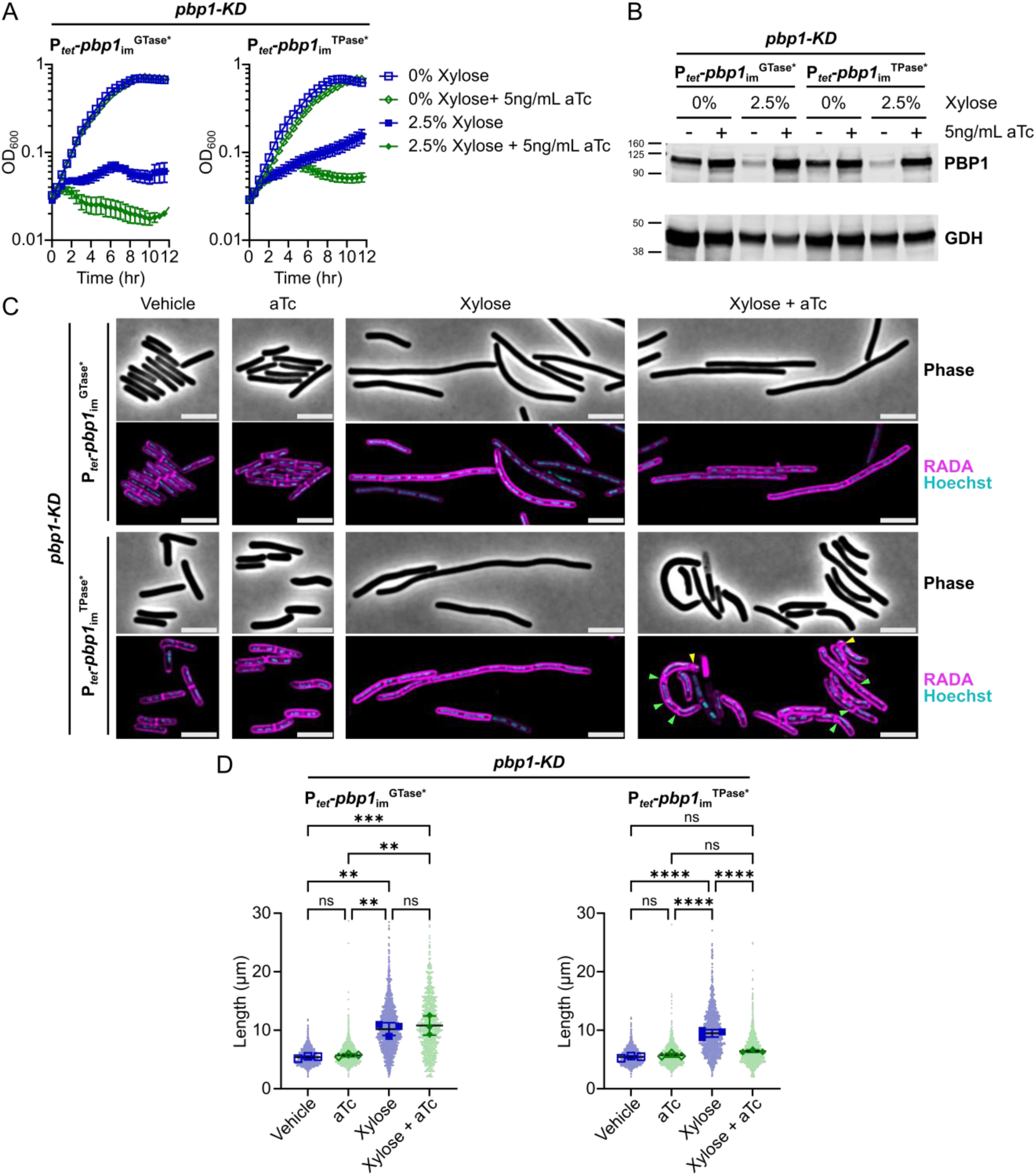
Both GTase and TPase activities of PBP1 are essential for *C. difficile* growth, but PBP1 TPase-deficient cells can still form septa and complete cell division. (A) Growth of *C. difficile* strains containing the xylose-inducible *pbp1*-KD cassette and aTc-inducible *pbp1*_im_^GTase*^ or *pbp1*_im_^TPase*^ complementation constructs was monitored by OD_600_. Mean and standard error were calculated across nine biological replicates. (B) Western blot analyses of PBP1 in cells exposed to vehicle or 2.5% xylose and/or 5 ng/mL aTc per the scheme in Fig. 2C. GDH (46.0 kDa) was used as a load control. Blots are representative of three independent experiments, and quantification across experiments is presented in Supplementary Fig. S6. (C) Fluorescence microscopy analyses of cells exposed to 2.5% xylose and/or 5 ng/mL aTc per the scheme in Fig. 2C. RADA labeling was used to visualize *de novo* peptidoglycan synthesis and Hoechst staining to visualize DNA. Images are representative of three independent experiments. Yellow arrows point to polar septa being formed, and green arrows point to instances of aberrant puncta near septa, bulky septa, and/or multi-septate cells. Scale bars represent 5 μm. (D) Quantification of cell lengths for >800 cells across three independent experiments. Dots indicate individual cells, and the larger, outlined symbols represent the mean cell length from each replicate. The mean and standard deviation were calculated across replicates; statistical significance was determined between replicates by a one-way ANOVA with Tukey’s multiple comparisons test. ns, not significant; ** p<0.01, *** p<0.001, **** p<0.0001.

### PBP1 TPase-deficient cells complete cell division but exhibit aberrant PG incorporation and uncontrolled septum synthesis

We next examined the morphology of conditional expression strains where PBP1’s GTase or TPase activities were genetically inactivated. Consistent with prior analyses showing that the chemical inhibition of PBP1’s GTase activity with MoeA inhibits cell division [12], *C. difficile* cells conditionally expressing *pbp1*_im_^GTase*^ exhibited a filamentous morphology and thus decreased septum synthesis (**Fig. 6C-D**). In contrast, *C. difficile* cells conditionally expressing *pbp1* ^TPase*^ had an overall cell length similar to vehicle-treated control cells, indicating that PBP1 transpeptidase activity is not strictly required for completing cell division. However, cell wall synthesis in *pbp1* ^TPase*^ cells appears dysregulated because greater variability in RADA labeling (puncta) and cell length was observed (**Fig. 6C**). The morphology of TPase-deficient cells was also altered, with curved and misshapen cells frequently forming. Furthermore, defects in septum placement near the cell poles, number, or thickness (bulges of RADA incorporation) were frequently seen (**Fig. 6C**), in contrast with vehicle-treated or *pbp1*_im_-expressing controls (**Fig. 3B**). Taken together, our data indicate that *C. difficile* cells conditionally expressing *pbp1* ^TPase*^ complete septum synthesis in an uncontrolled manner, generating irregular septa with abnormal RADA labeling.

### PBP3 promotes aberrant PG incorporation and abnormal septum synthesis in PBP1 TPase-deficient cells in a non-catalytic manner

Our finding that *pbp1*_im_^TPase*^ cells still efficiently incorporate the RADA D-amino acid analog, which can only be incorporated into peptidoglycan through the activity of a TPase [63], indicates that another enzyme with TPase activity must be mediating RADA incorporation. *C. difficile* encodes three bPBPs with TPase activity: the sporulation-specific bPBP, SpoVD, which pairs with the SEDS enzyme SpoVE [12,64] and is essential for sporulation; the sidewall-synthesis enzyme PBP2, which pairs with the SEDS enzyme RodA [12]; and an orphan bPBP, PBP3, which is induced during sporulation but has only subtle effects on sporulation and vegetative growth [12,65]. Since SpoVD functions exclusively during sporulation and is undetectable in vegetative cells [12,64] and PBP2 appears to function exclusively during cell elongation [12], we hypothesized that PBP3 could contribute to the aberrant peptidoglycan synthesis observed in cells conditionally expressing *pbp1*_im_^TPase*^. This hypothesis was based on the following observations: (i) PBP3 promotes asymmetric division when SpoVD catalytic activity is inactivated [65] and (ii) loss of *pbp3* leads to a statistically significant increase in *C. difficile* cell length (∼25%) during vegetative growth [12], implying that PBP3 is present at low levels during vegetative growth and may modulate vegetative cell division in *C. difficile*.

To test whether PBP3 contributes to abnormal peptidoglycan incorporation observed in PBP1 TPase-deficient cells, we conditionally expressed the *pbp1*_im_^TPase*^ construct in a mutant lacking *pbp3*. Remarkably, conditionally expressing the *pbp1*_im_^TPase*^ construct in the absence of PBP3 induced severe filamentation and decreased overall RADA incorporation, similar to cells depleted of PBP1 (*pbp1*-KDΔ*pbp3*/P*_tet_*-*pbp1*_im_^TPase*^ vs. *pbp1*-KD/P*_tet_*-*pbp1*_im_^TPase*^, **Fig. 7**). These data indicate that the aberrant peptidoglycan insertion and uncontrolled formation of septa observed in PBP1^TPase*^-producing cells relies on the presence of PBP3. Consistent with this conclusion, complementing back the expression of *pbp3* in the *pbp1*-KDΔ*pbp3*/P*_tet_*-*pbp1* ^TPase^ strain prevented filament formation and restored the cell length phenotype of the WT background conditionally expressing *pbp1* ^TPase*^; it also prevented the uncontrolled septum formation and aberrant RADA incorporation that distinguishes this strain (**Fig. 7**). Notably, western blot analyses confirmed that PBP1^TPase*^ is stably expressed in the Δ*pbp3* background, indicating that PBP3 does not impact PBP1^TPase*^ stability but is instead required for triggering septum formation in the PBP1^TPase*^-producing cells (Supplemental Fig. S7).

**Figure 7:**
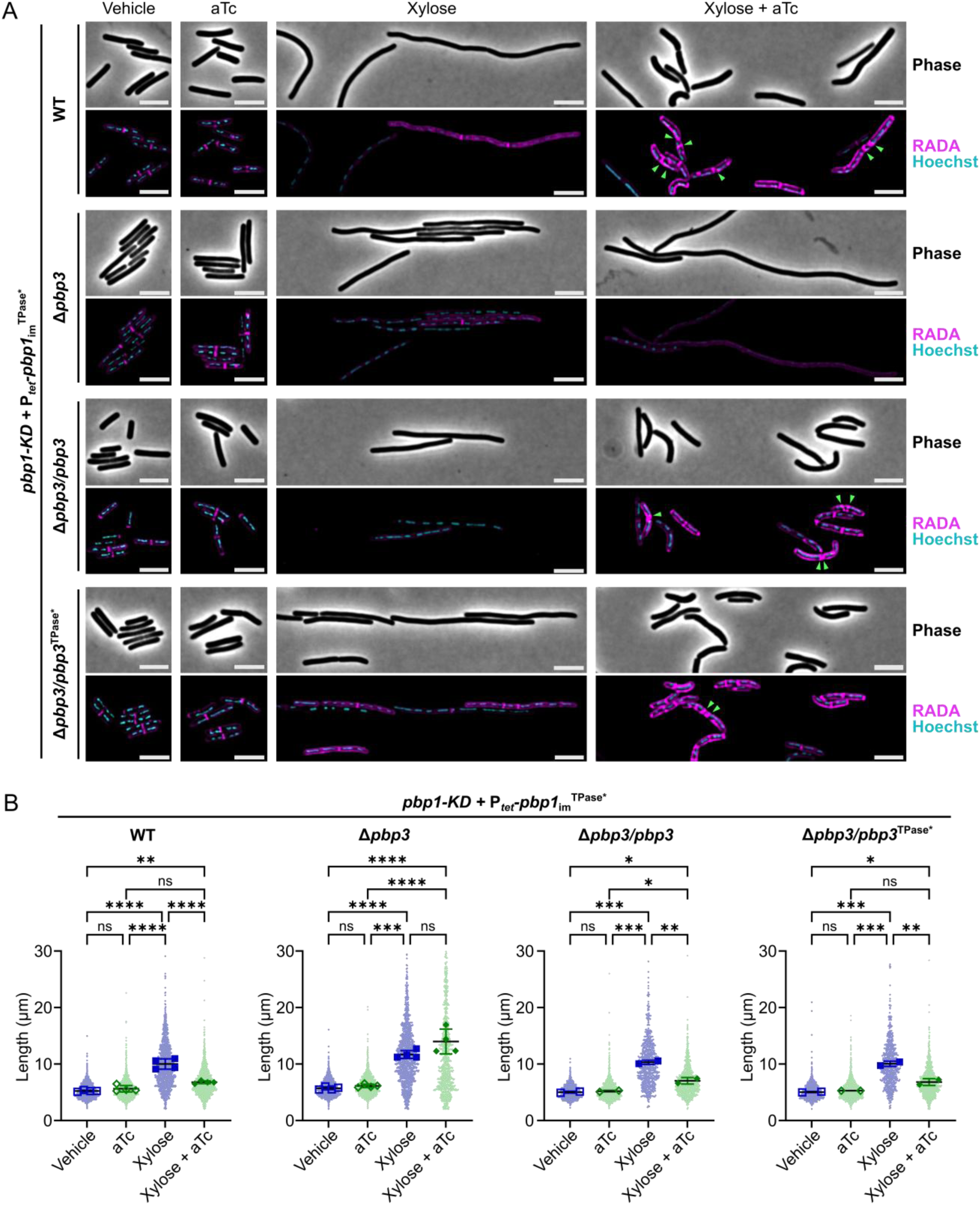
PBP3 promotes uncontrolled septum synthesis and aberrant peptidoglycan incorporation in PBP1 TPase-deficient cells. (A) Growth of *C. difficile* strains containing the xylose-inducible *pbp1*-KD cassette in WT or a Δ*pbp3* mutant and a plasmid-encoded, aTc-inducible *pbp1*_im_^TPase*^ complementation construct. Complementation of Δ*pbp3* was accomplished by expressing *pbp3* or *pbp3*^TPase*^ as a bicistronic construct with *pbp1*_im_^TPase*^ under the P*_tet_* promoter. Cells exposed to 2.5% xylose and/or 5 ng/mL aTc per the scheme in Fig. 2C were subject to RADA labeling for 10 min to visualize *de novo* peptidoglycan synthesis and Hoechst staining for DNA. Images are representative of 2-4 independent experiments. Green arrows indicate examples of irregular septa being formed. Scale bars represent 5 μm. (B) Quantification of cell lengths for >600 cells across 2-4 independent experiments. Dots indicate individual cells, and the larger, outlined symbols represent the mean cell length from each replicate. The mean and standard deviation were calculated across replicates; statistical significance was determined between replicates by a one-way ANOVA with Tukey’s multiple comparisons test. ns, not significant; ** p<0.01, *** p<0.001, **** p<0.0001.

Since functionally redundant bPBPs can compensate for the loss of the catalytic activity of the FtsI bPBP in some bacteria [3,59], we tested whether PBP3’s TPase activity was enabling the *pbp1* ^TPase*^ strain to complete cell division and incorporate RADA by complementing the Δ*pbp3* strain conditionally expressing *pbp1* ^TPase^ (*pbp1*-KDΔ*pbp3*/P_tet_-*pbp1* ^TPase^) with a construct encoding a TPase catalytic mutant of PBP3. This construct encodes a single amino acid substitution in the conserved catalytic SxxK motif of PBP3’s active site, S299A (*pbp3*^TPase*^). Surprisingly, the *pbp3*^TPase*^ complementation construct was also sufficient to restore normal septum formation and RADA incorporation in Δ*pbp3* cells conditionally expressing the *pbp1* ^TPase*^ allele (**Fig. 7**). Thus, PBP3 is not the TPase responsible for incorporating RADA at the deformed septa and peptidoglycan bulges observed in cells conditionally producing PBP1^TPase*^, but the presence of PBP3 is needed to induce uncontrolled septum synthesis in PBP1 TPase-deficient cells.

### PBP3 is a component of the *C. difficile* divisome complex

Since a possible explanation for these data is that PBP3 plays a regulatory role in activating septum synthesis, perhaps by enhancing the GTase activity of PBP1^TPase*^ and ultimately causing dysregulation of peptidoglycan synthesis in these cells, we tested whether PBP3 is recruited to the vegetative *C. difficile* divisome. Specifically, we compared the localization profile of PBP3 to those of known divisome proteins, PBP1 and FtsZ [12,66], using mSc-I3 protein fusions. Constructs encoding the fusion proteins inducibly expressed from the P*_tet_* promoter were integrated downstream of the *pyrE* locus in WT *C. difficile*. After transiently inducing the expression of the fusion protein constructs with aTc and visualizing *de novo* peptidoglycan synthesis with HADA, we found that PBP3 localized both to septa and the sidewall, similar to PBP1 (**Fig. 8A**). In demographs comparing mSc-I3 and HADA labeling across the medial axis of cells, PBP1 and PBP3 recruitment to mid-cell is coincident with the onset of septum synthesis, as indicated by HADA incorporation at mid-cell (**Fig. 8B**). By comparison, FtsZ localization was restricted to the mid-cell, and this localization was observed prior to septum synthesis (**Fig. 8A-B**). These data indicate that PBP3 can localize to the site of septum synthesis and the sidewall, similar to PBP1, and therefore may play a role both in vegetative cell division as part of the *C. difficile* divisome as well as in sidewall synthesis.

**Figure 8:**
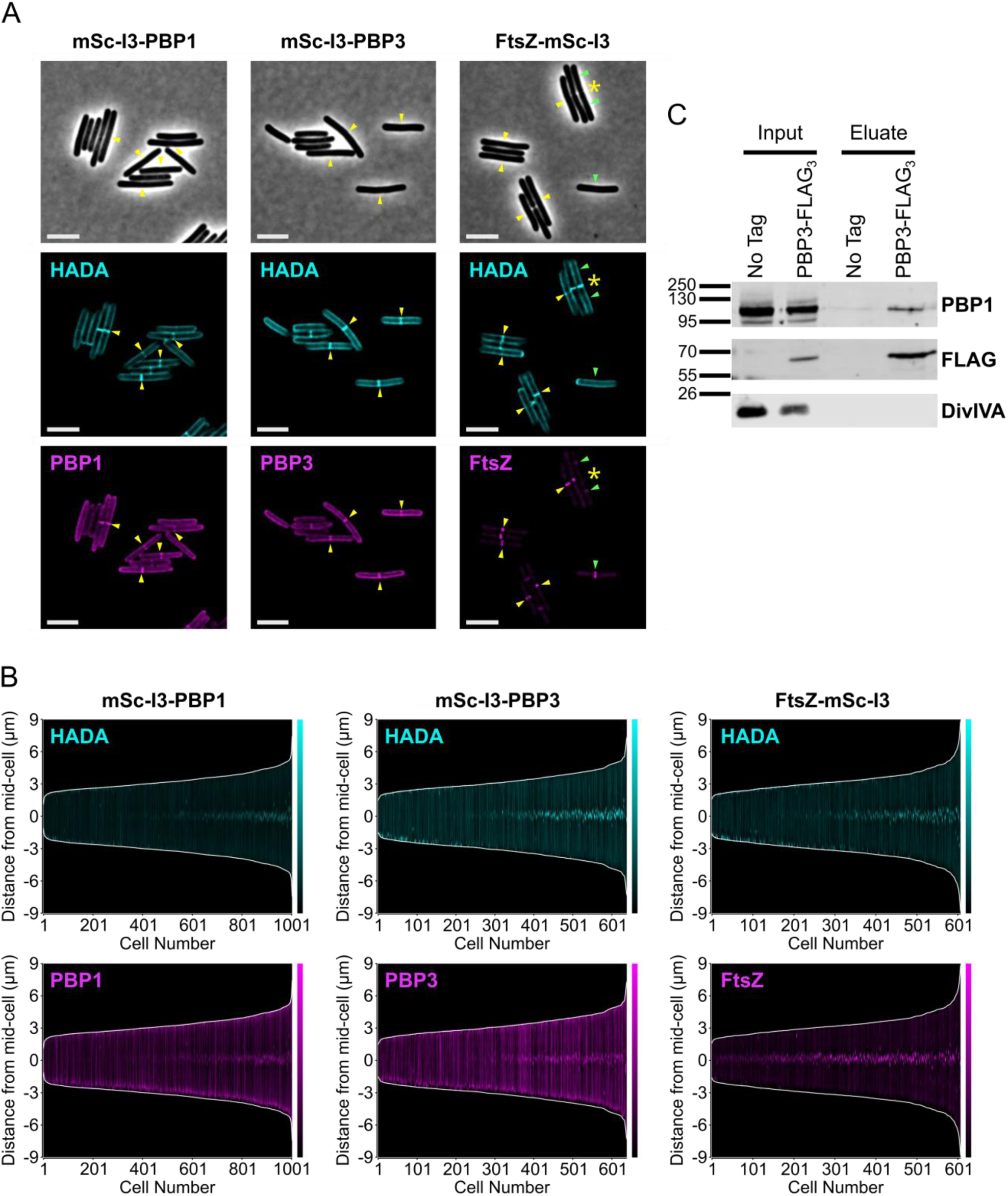
PBP3 can localize to the *C. difficile* divisome and forms a complex with PBP1. (A) Localization of fluorescently-tagged proteins in *C. difficile* carrying either an N-terminal (PBP1 and PBP3) or C-terminal (FtsZ) mSc-I3 tag under the control of an aTc-inducible promoter. Cells were cultured in the presence of 5 ng/mL aTc for 1 hr, exposed to HADA for 10 min to label *de novo* peptidoglycan synthesis, and then fixed for microscopy. Yellow arrows indicate co-localization of mSc-I3 signal and HADA. For FtsZ-mSc-I3, septa with no mSc-I3 signal are indicated with a yellow asterisk (*); these represent late-stage septa with no FtsZ localization. Green arrows indicate that the mSc-I3 signal does not co-localize with noticeable HADA incorporation; these likely represent early FtsZ-rings that have not yet initiated septum synthesis. Scale bars represent 5 μm. (B) Demographs depicting the medial axis fluorescence profile of >600 cells analyzed via MicrobeJ. Each cell is indicated on the x-axis, and cells are ordered by length. The cell length is indicated by the white lines, whereas the HADA or mSc-I3 signal is depicted in cyan or magenta, respectively. The HADA signal intensity was normalized across all images and demographs to facilitate direct comparisons between samples, whereas the mSc-I3 signal was scaled independently for each individual reporter construct. (C) Co-immunoprecipitation analysis of PBP3-FLAG_3_. Δ*pbp3* strains were engineered to harbor a P*_xyl_*-*pbp3*-*FLAG*_3_ construct or a *P_xyl_*-*pbp3* no-tag control. Cultures were grown in the presence of 0.5% xylose, then cells were lysed, membrane proteins were solubilized with 0.5% DDM, and anti-FLAG resin was used to immunoprecipitate the bait protein. PBP3-FLAG_3_, PBP1, and the negative control DivIVA were detected by western blotting.

We next assessed whether PBP3 interacts with PBP1 during vegetative cell growth. Notably, we previously reported that *C. difficile* PBP1 and PBP3 interact in a bacterial two-hybrid assay performed in *E. coli* and this interaction is also observed during sporulation [65]. To determine if PBP1 and PBP3 also interact during vegetative growth in *C. difficile*, we generated strains that inducibly express constructs encoding either PBP3 or PBP3-FLAG_3_ (prior RNA-Seq analyses indicated that *pbp3* is expressed at low levels in vegetative cells [67]). Actively growing cultures of the resulting strains were treated with formaldehyde crosslinker to stabilize protein-protein interactions, lysed, and the membrane proteins were solubilized with n-dodecyl-β-D-maltoside before performing the anti-FLAG co-immunoprecipitation. PBP3-FLAG_3_ co-immunoprecipitated with PBP1 but not with the negative control DivIVA, a membrane-associated protein that is also present at the site of division [68], suggesting that PBP3 and PBP1 are able to interact either directly or indirectly in vegetatively growing *C. difficile* (**Fig. 8C**). Collectively, our findings indicate that PBP3 and PBP1 localize to the site of septum synthesis in dividing vegetative *C. difficile* and that these two proteins interact, directly or indirectly, as part of the same complex. Based on these observations, we conclude that PBP3 is an accessory component of the *C. difficile* divisome complex that functions to promote its septum synthesis activity.

## Discussion

By taking advantage of newly developed genetic tools in *C. difficile* to establish a CRISPRi-based trans-complementation system, we performed structure-function analyses of PBP1, the essential peptidoglycan synthase required for cell growth and division [51]. This system allowed us to rapidly determine the requirement for discrete domains within PBP1 and its enzymatic activities, which would have been difficult to accomplish by engineering mutations in the native *pbp1* locus within the *C. difficile* genome.

Our analyses revealed that PBP1’s N- and C-terminal IDRs are both dispensable for bacterial growth and division. This result was surprising because the extracellular IDR of *B. subtilis* PBP1 is important for normal growth of *B. subtilis* likely by helping PBP1 detect gaps in the cell wall to repair [39]. While we observed slight morphological abnormalities in the PBP1^ΔCT^- and PBP1^ΔNTΔCT^-expressing *C. difficile* that could be consistent with uneven deposition of peptidoglycan on the sidewall [39,69], the increase in cell curvature in these cells was not statistically significant (Supplementary Fig. S3). Since our studies of PBP1 IDR function were carried out in standard laboratory conditions, it is possible that the *C. difficile* PBP1 IDRs become important for bacterial growth or cell morphology in non-standard laboratory conditions such as upon cell envelope stress. Indeed, in *B. subtilis*, a *pbp1* IDR mutant exhibits a more severe phenotype in a sensitized mutant background lacking GpsB and the other three *B. subtilis* aPBPs [39]. It is also possible that the residual WT PBP1 protein present during *pbp1*-KD seen in western blot analyses could obscure the functional requirement of PBP1’s IDRs in *C. difficile* (**Fig. 2D**)(Supplementary Fig. S1A), so analyzing IDR truncation mutations in the native *pbp1* locus in various stress-inducing conditions could provide insight into these questions.

In contrast with the dispensability of the IDRs for PBP1 function, we determined that the GI domain, a novel structural feature in *C. difficile* PBP1 conserved in only a subset of bacterial families, is critical for PBP1 function in *C. difficile*. The location of the GI domain is reminiscent of the UB2H domain in the division-associated aPBP in *E. coli*, PBP1b, which serves as a critical regulatory domain for interactions with essential PBP1b regulators such as LpoB and CpoB [31–34,70]. While the precise function of the GI domain remains unknown, we showed that PBP1^ΔGI^ is stably produced and has a similar localization profile as WT PBP1. It is possible that the GI domain either allosterically activates the GTase or TPase activity of PBP1 directly or serves as an interaction site for as-yet unidentified PBP1 regulators. The conservation of the GI domain in aPBPs from multiple bacterial families indicates that the GI domain may govern aPBP function in diverse bacteria found in anaerobic environments ranging from the human body to deep-sea hydrothermal vents [45,46].

We further demonstrated that PBP1’s GTase activity is essential for PBP1 function during cell growth and division in *C. difficile* (**Fig 6**), consistent with prior analyses using the aPBP GTase inhibitor Moenomycin A [12]. In contrast, we found that PBP1’s TPase activity is essential for PBP1 to support cell growth but is dispensable for its function during cell division (**Fig 6**). This finding was surprising on two levels. First, a prior chemical screen using β-lactam antibiotics predicted that PBP1’s TPase activity is likely dispensable for *C. difficile* growth because β-lactams that potently inhibit PBP1 TPase activity *in vitro* do not inhibit *C. difficile* growth [58]. Notably, our genetic interrogation of PBP1’s TPase activity establishes that PBP1 TPase activity is essential for growth. However, it also reveals that PBP1 TPase activity is dispensable for *C. difficile* cell division, although PBP1 TPase*-producing cells exhibit morphological defects and dysregulated septal and sidewall peptidoglycan synthesis (**Fig 6**). Second, we found that the uncontrolled septum synthesis observed in PBP1 TPase-deficient cells depends on the presence, but not the activity, of the non-essential enzyme PBP3 (**Fig 7**). The non-enzymatic role that we uncovered for PBP3 in promoting dysregulated septal and sidewall peptidoglycan synthesis in PBP1 TPase-deficient cells was somewhat unexpected because functionally redundant TPases have been shown to supply the TPase activity needed for essential SEDS-bPBP complexes to mediate cell growth in other Gram-positive bacteria [3,59].

These findings lead to the question of which enzyme(s) supply the TPase activity that mediates septum synthesis and dysregulated RADA incorporation in PBP1^TPase*^/PBP3^TPase*^ cells (**Fig 7**). One possibility is that since the CRISPRi-KD of *pbp1* is not complete, the residual WT PBP1 present in these cells may contribute to the RADA incorporation at these sites. The elongation-specific bPBP, PBP2, may also contribute to dysregulated septal PG synthesis, but this seems less likely given the clear separation in cell elongation and division roles in *C. difficile* and other organisms [12,71–74]. Alternatively, the TPase activity supplied at these sites of dysregulated peptidoglycan synthesis may be provided by another class of enzyme: the LDTs, since *C. difficile* encodes five LDTs and requires at least one for viability [61,62,75,76]. The LDTs could be responsible for the formation of aberrant septa and peptidoglycan bulges observed in PBP1^TPase*^/PBP3^TPase*^ cells, perhaps by interacting with PBP1 or PBP3.

Regardless, our analyses provide novel insight into the role of PBP3 during vegetative growth in *C. difficile*. While we previously showed that PBP3 is a non-essential, sporulation-induced bPBP that promotes asymmetric division [65], the current study revealed that PBP3 functions to promote septum synthesis during vegetative growth because PBP1 TPase-deficient cells filament in the absence of PBP3 (**Fig 7**). While this cryptic function was uncovered in a PBP1 TPase-deficient strain, it is consistent with our prior finding that Δ*pbp3* cells are slightly longer than WT *C. difficile* during vegetative growth [12]. Thus, PBP3 appears to specifically modulate septal peptidoglycan synthesis by PBP1 during both asymmetric and vegetative cell division.

Consistent with this interpretation, we previously showed that PBP3 interacts directly with PBP1 when expressed in *E. coli* in a bacterial two-hybrid assay and that PBP1 co-immunoprecipitates with PBP3 during sporulation [65]. In this study, we found that PBP1 also co-immunoprecipitates with PBP3 during vegetative growth (**Fig. 8C**) and localizes to vegetative septa (**Fig. 8**). Similar interactions between bPBPs and aPBPs have been reported in γ-Proteobacteria to promote divisome and elongasome function [37,77,78]. For instance, the *Acinetobacter baumanii* aPBP PBP1A is part of a complex containing the division-associated bPBP, FtsI, and this complex promotes septum synthesis [78]. However, it should be noted that *A. baumanii* differs from *C. difficile* because its aPBP activity alone cannot drive septum synthesis [78]. Additionally, the division-associated *E. coli* aPBP, PBP1b, interacts directly with FtsI, leading to its recruitment to the mid-cell during division [37], while the elongasome-associated bPBP in *E. coli*, PBP2, directly stimulates PBP1a GTase activity, and the combination of these bPBP and aPBP enzymes cooperatively enhance their TPase activity [77]. Therefore, bPBPs can directly stimulate aPBP enzymatic activity. Given these findings in *E. coli*, it is possible that *C. difficile* PBP3 directly stimulates the enzymatic activity of PBP1 such that PBP3 induces uncontrolled GTase activity in PBP1^TPase*^ cells, leading to dysregulated peptidoglycan synthesis and aberrant septa formation. Future investigation into the cross-talk between PBP1 and PBP3 in *C. difficile* will generate new insight into the interplay between PBP enzymes in bacteria.

## MATERIALS AND METHODS

### Bacterial strains and growth conditions

All *C. difficile* strains are derived from the 630Δ*erm* strain background and listed in Supplementary Table S1. Chromosomally-encoded mutations were generated in a Δ*pyrE* strain using *pyrE*-based allele coupled exchange as previously described [52]. *C. difficile* was cultured in brain heart infusion medium supplemented with 0.5% yeast extract and 0.1% L-cysteine (BHIS) with thiamphenicol (5 μg/mL) to maintain episomal plasmids, and/or kanamycin (50 μg/mL) and cefoxitin (8 μg/mL) as needed for genetic manipulation. *C. difficile* defined medium (CDDM) [79] was used for allele-coupled exchange to select for *pyrE* restoration. *C. difficile* cultures were grown at 37°C in an anaerobic chamber using a gas mixture containing 85% N_2_, 5% CO_2_, and 10% H_2_.

Plasmids used in this study were cloned using Gibson assembly, maintained in *E. coli* DH5α, and confirmed by Sanger or nanopore sequencing. *E. coli* strains containing plasmids used in this study are listed in Supplementary Table S2, with links to plasmid maps containing primer sequences used for cloning. To introduce constructs into *C. difficile*, plasmids were first transformed into *E. coli* HB101/pRK24 and then conjugated into *C. difficile* as previously described [80]. *E. coli* cultures were grown in LB at 37°C supplemented as needed with chloramphenicol (20 μg/mL), ampicillin (100 μg/mL), or kanamycin (30 μg/mL).

### Conditional expression of *pbp1* and growth curve analysis

*C. difficile* starter cultures were grown for ∼2 hr, back-diluted 1:20 into BHIS with 0% or 2.5% xylose, and grown for an additional 2 hr to pre-deplete PBP1 using the xylose-inducible CRISPRi system. Cultures were then adjusted to an OD_600_ of 0.05 in media containing 0% or 2.5% xylose and/or 5 ng/mL aTc to induce the *pbp1* complementation constructs, immediately added to a 96-well plate, and the OD_600_ was measured every 30 minutes in an Epoch 2 plate reader (BioTek) in an anaerobic chamber. The xylose and aTc concentrations were titrated to identify the optimal concentrations of xylose to efficiently knock-down endogenous *pbp1* expression and aTc to allow for trans-complementation. For fluorescence microscopy and western blot analysis, the cultures were allowed to grow an additional 4 hr before samples were taken. The concentration of vehicle (water for xylose and ethanol for aTc) were normalized across all samples.

### Fluorescent probes

The fluorescent D-amino acids RADA or HADA (Tocris) were used to label *de novo* peptidoglycan synthesis. Hoechst 33342 (Molecular Probes) was used to stain bacterial DNA. 500 μL of *C. difficile* culture were exposed to 50 μM RADA/HADA and/or 20 μg/mL Hoechst for 10 min, then fixed with a mixture of 100 μL 16% paraformaldehyde and 20 μL 1 M NaPO_4_ buffer (pH 7.4) for 30 min at room temperature and 30 min on ice [81]. Fixed cells were then washed three times with 1 mL 1XPBS before imaging.

### Protein localization with mScarlet-I3 fusions

*C. difficile* strains were engineered to express chromosomally-encoded P*_tet_*-*mScarlet-I3-pbp1*, P*_tet_*-*mScarlet-I3-pbp3*, or P*_tet_*-*ftsZ*-*mScarlet-I3* constructs downstream of *pyrE*. Strains were cultured to logarithmic phase in BHIS, then exposed to 5 ng/mL aTc (for *pbp1* and *pbp3* constructs) or 0.5 ng/mL aTc (for *ftsZ* construct) for 1 hr. Induced cultures were labeled with HADA and fixed as described above. After cell fixation, cells were incubated overnight at room temperature in the dark to allow chromophore maturation prior to imaging, as previously described [81].

### Fluorescence microscopy

Microscopy samples were imaged on agarose pads (1% agarose in 1XPBS). Phase-contrast and fluorescence micrographs were acquired with a Leica DMi8 inverted microscope equipped with a 63X 1.4 NA Plan Apochromat oil-immersion phase-contrast objective, a high precision motorized stage (Pecon), and in a 37°C incubator (Pecon). Excitation light was generated by a Lumencor Spectra-X multi-LED light source with integrated excitation filters. An XLED-QP quadruple-band dichroic beam-splitter (Leica) was used (transmission: 415, 470, 570, and 660 nm) with an external filter wheel for all fluorescent channels. HADA and Hoechst were excited at 395/25, and emitted light was filtered using a 440/40-nm emission filter, with 120 ms (HADA) or 100 ms (Hoechst) exposure times; RADA and mScarlet-I3 were excited at 550/28 nm, and emitted light was filtered using a 590/50-nm emission filter, with 300 ms (RADA) or 150 ms (mScarlet-I3) exposure times. Light was detected using a Leica DFC 9000 GTC sCMOS camera. 1-2 μm z-stacks were taken with 0.21 μm z-slices. Images were acquired using the LASX software, and fluorescence images were deconvolved using Leica Small Volume Computational Clearing with the following settings: refractive index 1.33, strength 60%, and regularization 0.05.

Images were processed using FIJI to select the best-focused z-plane for each channel and adjust the image brightness and contrast of images. Identical minimum and maximum values were applied to a given channel across all images in a figure panel so that direct comparisons can be made across images within a figure panel, unless otherwise specified in the figure legend. Demographs were generated with MicrobeJ in FIJI, using segmentation masks generated by SuperSegger [82].

### Quantification of cell length and width

To quantify the length and width of cells, phase-contrast images were segmented using the MATLAB-based image analysis pipeline SuperSegger [82]. To enable analysis of filamentous cells, the default “60xec” settings were modified with the following: CONST.superSeggerOpti.MAX_WIDTH = 1e10; and CONST.seg.OPTI_FLAG = false. Cells cut off by the edge of the image were excluded from the analysis and the minimum cell length was set to 2 μm to avoid inadvertently including cellular debris in the dataset.

### Western blot analysis

*C. difficile* cultures were pelleted, resuspended in 25 μL 1XPBS, freeze-thawed three times, mixed with 25 μL EBB buffer (9 M urea, 2 M thiourea, 4% SDS, 2 mM β-mercaptoethanol), and boiled for 20 min to lyse cells. To normalize the protein load across samples, the lysates were diluted based on the starting OD_600_ of the sample prior to lysis, and then the same volume of lysate was run using SDS-polyacrylamide gel electrophoresis (SDS-PAGE) on a 10% polyacrylamide gel. Proteins were transferred to polyvinylidene difluoride membranes, which were subsequently probed with primary antibodies: rabbit polyclonal anti-PBP1 TF134 [65] at 1:1,000 dilution; rabbit polyclonal anti-DivIVA TF142 (this study) at 1:1,000 dilution; chicken polyclonal anti-GDH (aCdGDH; Thermo) at 1:10,000 dilution; and/or a mouse monoclonal M2 anti-FLAG antibody (Sigma) at 1:5,000. Anti-rabbit, anti-chicken, and anti-mouse IR800 or IR680 secondary antibodies (LI-COR Biosciences, 1:20,000) were used to detect bands with a LI-COR Odyssey CLx imaging system. Quantification of westerns was performed using LI-COR imaging software Image Studio^TM^ Lite with background subtraction. Band intensities were normalized to GDH for each sample, and further normalized to the vehicle-treated control to calculate the fold-change in PBP1 levels relative to the control cells.

### Antibody production

The anti-DivIVA antibody used for western blots in this study were raised in rabbits by Cocalico Biologicals against *C. difficile* DivIVA-His_6_ purified from *E. coli*. DivIVA-His_6_ was produced in BL21(DE3) *E. coli* harboring pET28a-*divIVA*-*His_6_* and purified by Ni^2+^-affinity purification as previously described [83]. Proteins were purified further by size exclusion chromatography (SEC) in a buffer containing 200 mM NaCl, 10 mM Tris-HCl pH 7.5, 5% glycerol, and 1 mM DTT, using a Superdex 200 Increase 10/300 GL (GE Healthcare) column. SEC fractions containing DivIVA, as determined by SDS-PAGE and western blotting with an anti-His_6_ antibody (HIS.H8; Thermo), were combined, concentrated, and used to raise antibodies. Antisera reactivity and specificity for DivIVA was validated by western blot against *C. difficile* lysate from WT and a *divIVA* CRISPRi knock-down strain.

### Co-immunoprecipitation (Co-IP) analysis

To immunoprecipitate FLAG-tagged proteins from vegetatively growing *C. difficile*, 200 μL of exponentially growing cultures of *C. difficile* harboring a xylose-inducible P*_xyl_*-*pbp3*-FLAG_3_ or a P*_xyl_*-*pbp3* un-tagged expression construct engineered downstream of the *pyrE* locus in the chromosome were spread on BHIS agar media containing 0.5% xylose and grown for 9 hr. Cells were scraped from 3 plates per strain and pooled in 750 μL of BHIS. A portion of cells was visualized by microscopy with RADA labeling to confirm that cells were actively dividing and forming septa. Co-immunoprecipitation was performed similar to previously described methods. Crosslinking of cells was performed with 0.25% final concentration of PFA for 15 minutes at 37°C followed by quenching with 350 mM glycine for 10 minutes on ice. Cells were then pelleted, re-suspended in 750 µL of FLAG IP Buffer (150 mM NaCl, 50 mM Tris-HCl pH 7.5), transferred to screwcap tubes containing MP Biomedicals Lysing Matrix E, and frozen at −80°C. Cell pellets were thawed and bead beat on an MP Biomedicals FastPrep-24 four times at 5.5 M/second for 1 minute for a total of four rounds of lysis, with 5 minutes on ice between rounds of bead-beating. Next, 1X HALT protease inhibitors and 0.5% final concentration of dodecyl-β-d-maltoside (DDM) detergent was added to lysed cells, and cell lysates were rotated at room temperature for 1 hour to solubilize membrane proteins. Lysates were clarified with centrifugation at 10,000 × g for 1 minute, and 200 µL of pre-equilibrated Anti-FLAG M2 magnetic resin was added to the clarified lysate. The slurry containing lysate and resin was rotated for 1 hour at room temperature. To remove unbound proteins, the resin was washed three times briefly and once for 15 minutes with 1 mL FLAG IP Buffer containing 0.5% DDM. This step was repeated a total of two times. Then, the resin was washed three times briefly and once for 15 minutes with FLAG IP Buffer containing no detergent. Finally, bound proteins were eluted from the Anti-FLAG resin with 100 μg/mL 3XFLAG peptide. Samples were boiled in 1X sample buffer (40% glycerol, 1M Tris pH 6.8, 20% β-mercaptoethanol, 8% SDS, and 0.04% bromophenol blue) for 10 minutes to reverse crosslinks and then analyzed by western blot.

## Acknowledgements

Thank you to Shailab Shrestha for helpful discussion and advice during the conception of this work. G.A.H. was supported by the National Institute of General Medical Sciences K12GM133314 through the Tufts Institutional Research Career and Academic Development Award Program, and this work was supported by National Institute of Allergy and Infectious Diseases R01AI122232 awarded to A.S.. Sponsors or funders did not play any role in the study design, data collection and analysis, decision to publish, or preparation of the manuscript.

